# Life history speed, population disappearances and noise-induced ratchet effects

**DOI:** 10.1101/2022.01.27.478031

**Authors:** Christopher J. Greyson-Gaito, Gabriel Gellner, Kevin S. McCann

## Abstract

Nature is replete with variation in the body sizes, reproductive output, and generation times of species that produce life history responses known to vary from small and fast to large and slow. Although researchers recognize that life history speed likely dictates fundamental processes in consumer-resource interactions like productivity and stability, theoretical work remains incomplete in this critical area. Here, we examine the role of life history speed on consumer-resource interactions by using a well used mathematical approach that manipulates the speed of the consumer’s growth rate in a consumer-resource interaction. Importantly, this approach holds the isocline geometry intact allowing us to assess the impacts of altered life history speed on stability (coefficient of variation, CV) without changing the underlying qualitative dynamics. Although slowing life history can be initially stabilizing, we find that in stochastic settings slowing ultimately drives highly destabilizing population disappearances, especially under reddened noise. Our results suggest that human-driven reddening of noise may decrease species stability because the autocorrelation of red noise enlarges the period and magnitude of perturbations, overwhelming a species’ natural compensatory responses via a ratchet-like effect. This ratchet-like effect then pushes species’ population dynamics far away from equilibria that can lead to precipitous local extinction.

## Introduction

Life history is a fundamental axis of variation within nature. Researchers have argued cogently that this variation tends to follow a “slow-fast” continuum where slow life histories have small population growth rates (low r), large body sizes, and long generation times while fast life histories have high population growth rates, small body sizes, and short generation times [1–4]. Indeed, empirical work has found remarkably consistent relationships that suggest body size is a key attribute of life history speed (Fig. 1a; [5]). These covarying traits along the “slow-fast” continuum can impact many ecological processes and properties including population growth, ecosystem productivity and stability [1,6]. Given the changing nature of abiotic variation under climate change [7,8] and the increasing propensity for fast life histories under global change [9,10], understanding how life history speed governs the ability for organisms and whole communities to persist (i.e., retain densities well above zero densities) is critical.

**Figure 1.**
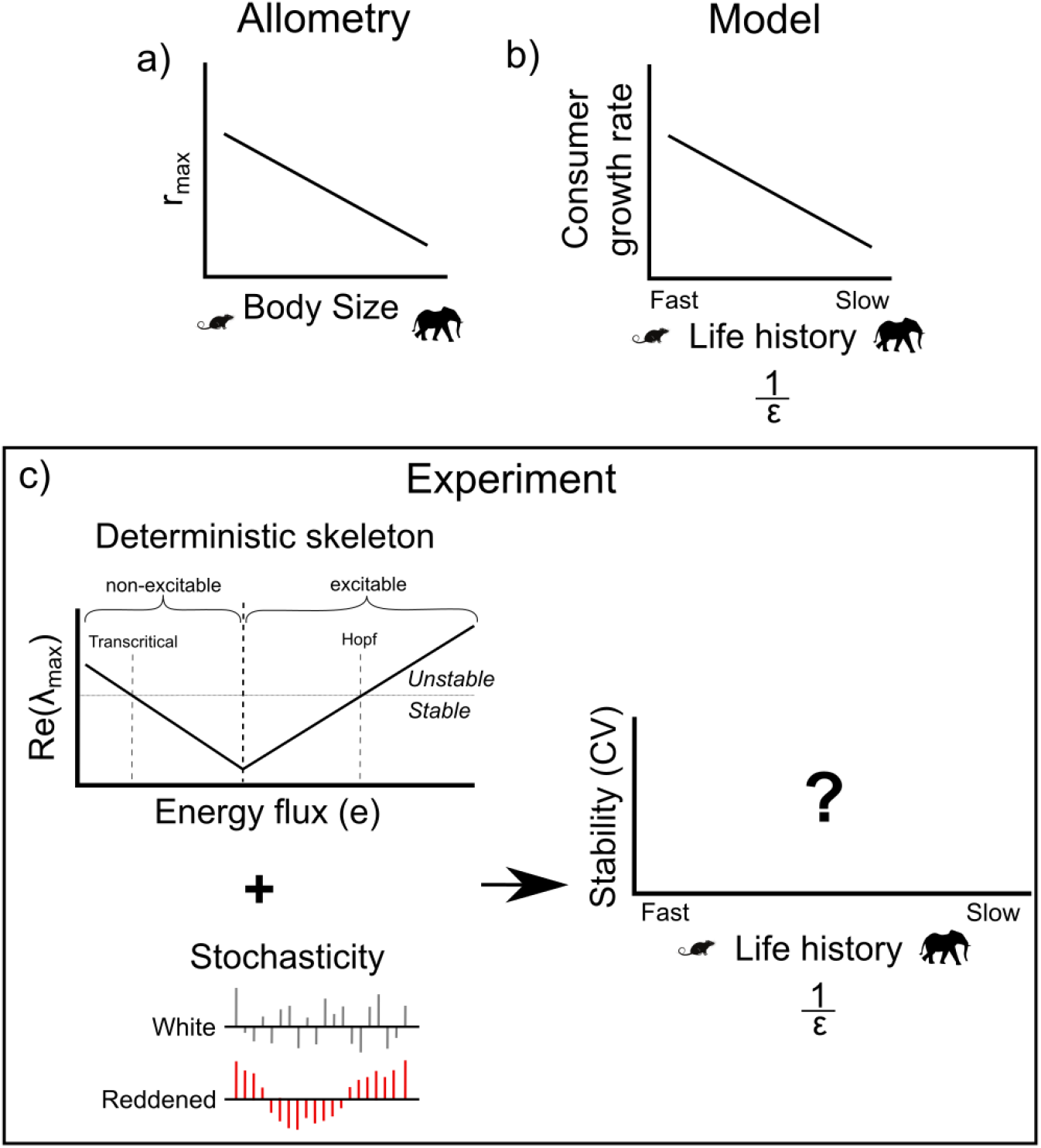
a) Empirical relationship between body size and population growth rate (r_max_) show the existence of slow-fast life histories (e.g., Savage et al. [1]). b) In our consumer-resource model, we use the parameter *ε* to scale the consumer’s growth rate to replicate the empirical relationship between body size and growth rate. *ε* scales the consumer equation such that the consumer growth rate decreases with increasing 1/*ε* producing a fast to slow life history continuum. This method simultaneously holds the isocline arrangement constant and so is an experiment that changes “life history speed” in and of itself. c) To explore the impacts of life history speed, we will examine how slowing the consumer’s life history through 1/*ε* impacts the stability of the C-R interaction (using coefficient of variation). Stochastic perturbations, from white to reddened, will be added to the consumer. Because we know that the underlying deterministic skeleton interacts with noise in different ways [21,22], we will set the efficiency parameter (*e*) of the C-R model to produce the non-excitable (i.e., real eigenvalues, monotonic dynamics) and excitable (i.e., complex eigenvalues, oscillatory decay dynamics) deterministic skeletons.

Theory has begun to unpack how population life history “speed” may tie into stability, with initial findings of fast and slow life histories amplifying or muting noise respectively. Specifically, research has found that because of the high population growth rates, faster life history organisms tend to produce over-compensatory dynamics (i.e., overshoot the equilibrium) and instability compared to slower organisms [11,12]. In contrast to fast organisms, slow organisms, with longer lives, buffer noise and thus maintain high stability. Consistent with this, empirical research has found negative correlations between population variability (one measure of stability) and both body size and generation time, and positive correlations between population variability and growth rates across multiple taxa and kingdoms [13–16]. Taken altogether, larger and slower organisms appear capable of buffering noise better than small and fast organisms.

Nonetheless, while large slow-growth organisms may be able to buffer perturbations, this same potentially stabilizing slow growth response can be destabilizing when perturbations are large enough to push population densities far away from the equilibrium [17,18]. Once a slow-growth species is pushed away from the equilibrium they become subject to the vagaries of different nonlinear dynamics (hereafter referred to as non-local nonlinear dynamics, see Box 1) that can fundamentally alter the outcome of the system including taking the dynamics close to zero or local extinction [17,18]. On the other hand, small fast-growing species may be less likely to be pushed far away from equilibrium because their rapid growth responses can keep them closer to equilibrium and local dynamic properties. These results suggest complex slow-fast life history stability responses such that some aspects of life history speed are beneficial (e.g., large and slow can buffer a perturbation) but also produces costs (e.g., large and slow growth can be detrimental once the population is pushed away from local equilibrium dynamics). Consequently, more research is required to understand how stability manifests along the slow-fast life history continuum.

Another related factor that will impact stability along the slow-fast continuum is the color of the stochastic noise (see Box 1 for definition). Climate change is known to be reddening environmental noise by increasing the spatial and temporal autocorrelation of climate variables [7,8] which then can act to lengthen periods of suboptimal conditions for organisms [19]. Under extended suboptimal conditions, populations can be driven to smaller and smaller numbers thus often hastening local extinction [19]. In a sense, these extended suboptimal conditions are akin to large perturbations (although each individual noise event is small). By pushing an organism’s dynamics far from equilibria, red noise is then also likely to cause non-local nonlinear dynamic outcomes. In this latter case, although not well explored, these autocorrelated perturbations could even lead beneficial perturbations to drive strong nonlinear effects that eventually produce local extinction (e.g., consumer increases in a consumer-resource interaction can lead to overshoot and dangerously low densities). These collective theoretical results suggest a complex range of dynamical responses that demand further understanding.

Towards understanding the complex responses of stochastic models, Higgins et al. [20] pointed out that the dynamics of an underlying deterministic model are critical to understanding the full stochastically forced model. Through elegantly decomposing a stochastic population model of Dungeness crab into a deterministic model (the deterministic skeleton, see Box 1) and a stochastic process, Higgins et al. [20] found that the novel responses produced from the interaction of the density-dependent dynamics and stochasticity matched empirical observations. Similarly, qualitatively different dynamics of the deterministic consumer-resource (C-R) model are known to yield qualitatively different stochastic dynamics. For example, weakly interacting deterministic C-R models are known to produce non-excitable monotonic dynamics (i.e., the deterministic equilibrium is stable with real eigenvalues) that when perturbed tend to simply produce noisy equilibrium dynamics [21,22]. As the stochastic dynamics suggest, the monotonic deterministic skeleton is aptly named non-excitable. In contrast, as the C-R interactions are strengthened, the deterministic equilibrium begins to show excitable dynamics with decaying cycles to the equilibrium (i.e., the deterministic equilibrium is stable with complex eigenvalues). Here, stochastic perturbations now tend to excite the underlying density dependent frequencies of the deterministic C-R skeleton producing stochastic cycles or quasi-cycles [22]. Again, the dynamics of the underlying deterministic skeleton is critical to understanding the full stochastic model.

Here, towards synthesizing slow-fast life history stability responses to perturbations, we examine how stability (i.e., defined as per many empirical studies as the coefficient of variation, CV) is influenced by life history speed along the slow-fast continuum. Specifically, we examine the C-R interaction – a fundamental building block of whole communities – because it has been well described by allometric arguments that are conducive to slow-fast theory [23,24]. We alter the consumer’s life history speed using a common mathematical scaler (*e*) employed for the analysis of slow-fast mathematical systems [25]. The parameter, *e*, allows us to mimic known variation in growth rates (Fig. 1a,b) and to alter the life history speed while maintaining the underlying qualitative dynamical conditions (i.e. the isocline geometry of the C-R model is preserved when changing *e* alone). We employ this model over a range of white to red noise perturbations (Fig. 1c). Finally, we vary the C-R interaction strength (i.e. energy flux via consumer conversion efficiency, *e*) to explore both the non-excitable and the excitable deterministic skeletons (Fig. 1c). In doing so, we cover a broad range in the underlying deterministic skeleton and seek a general answer to the role slow-fast life histories play in stability. Overall, we show that life history speed drives a range of stability responses to noise with slow life histories especially sensitive to non-local nonlinear dynamics and population disappearances when perturbed by reddened noise.

### Box 1 Key terms and definitions

#### Deterministic skeleton

The deterministic skeleton mathematically describes the processes of interest in the system being modeled. These are usually a dynamical system (e.g., ordinary differential equation or difference equations) and are effectively the model without stochastic processes [26].

#### Noise/Stochasticity

In ecology, noise is considered to be whatever we do not understand in the system [26,27]. Modelers add stochastic (random) noise to the deterministic skeleton. The interaction between the deterministic skeleton and noise can generate radically different dynamical responses, providing key insights into ecological phenomena (see review by Coulson et al. [27]).

#### Red & white noise

Noise from time points close together that are similar to each other (positive autocorrelation) is classified as red noise. Red noise is the opposite of blue noise where noise from time points close together are completely different (negatively autocorrelated). White noise lies between red and blue noise, that is, noise from time points close together are neither always similar nor always different (i.e. neither positively nor negatively autocorrelated).

#### Stability

Here, we define stability in terms of persistence where increasing the lower limit of population density away from zero increases stability/persistence (general stability in McCann [28]). Notably, the lower a population’s density is the greater the likelihood that a stochastic process can push it to local extinction. The coefficient of variation (CV) is a common theoretical and empirical metric that normalizes standard deviation relative to the mean and so is a good metric for general stability (see Figure 1c). A high SD relative to a low mean means that the population necessarily attains a very low density where risk of collapse is high.

#### Non-local nonlinear dynamics

Nonlinear dynamics occur in systems that react disproportionately to initial conditions or a small perturbation. These dynamics can include chaos and limit cycles. In this study, we differentiate between local and non-local nonlinear dynamics. Local nonlinear dynamics are dynamics when system trajectories are close to the equilibria in phase space. Non-local nonlinear dynamics are dynamics when system trajectories are far from equilibria.

#### Quasi-cycle

Quasi-cycles are a type of dynamic behaviour that occur when stochasticity resonates with damped oscillations surrounding an interior equilibrium [21,26,29]. Frequencies in the stochastic noise that most closely resemble the period of the damped oscillations are amplified. Thus, a power spectrum would show all frequencies with the frequency of the damped oscillations having the highest power [21].

#### Quasi-canard

Quasi-canards are another type of dynamic behaviour that occur when stochasticity induces a pattern similar to a deterministic canard. A deterministic canard occurs when a system’s solution follows an attracting state space area (manifold), passes over a critical point along this manifold, and then follows a repelling manifold [30]. In the C-R model, the canard solution slowly follows the resource isocline (the attracting manifold) until the maximum point of the isocline is reached (the critical point), then the solution quickly jumps to the consumer axis before slowly following the consumer axis (the repelling manifold). Finally, the solution quickly jumps back to the resource isocline and repeats the canard cycle. When stochasticity is introduced, trajectories combine both small oscillations around the equilibria and large relaxation oscillations qualitatively similar to deterministic canards. Generally, these patterns are called mixed mode oscillations [30]. However, we use the term quasi-canards in this study to evoke the importance of stochastic noise similar to the importance of stochastic noise in quasi-cycles.

## Methods

### Model

In this study, we used the classic Rosenzweig-MacArthur consumer-resource (C-R) model

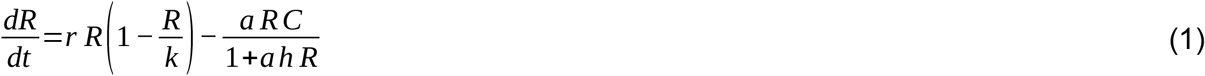

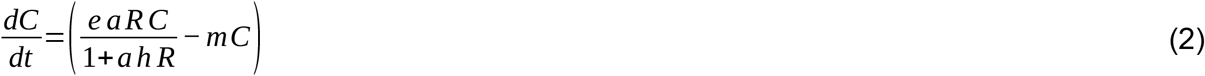

where *r* is the intrinsic growth rate of the resource (R), *k* is the carrying capacity of the resource, *a* is the attack rate of the consumer (C), *e* is the efficiency or conversion rate of consumed resources into new consumers, *h* is the handling time, and *m* is the consumer mortality.

Previous exploration has illustrated the multiple bifurcations and different dynamics of this model [14,31]. The bifurcations are dictated by the energy entering the consumer versus exiting the consumer (the energy flux of the consumer) or as an equation:

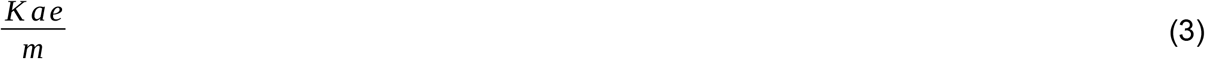

As the energy entering the consumer increases relative to the energy exiting the consumer (the numerator of (3) increases relative to the denominator), the first bifurcation is a transcritical bifurcation (switching of stable points) creating an interior stable equilibrium (see Figure 1c). At first, this equilibrium is monotonic but then exhibits damped oscillations. The next bifurcation is a Hopf bifurcation (the stable interior equilibrium becomes unstable) leading to period cycling. The mathematics underlying the shift from monotonic to dampened oscillations for the stable interior equilibrium is a shift in eigenvalues from real (no imaginary part) to complex (with an imaginary part). Gellner et al. [22] termed these two regions the non-excitable region and the excitable region respectively (see Figure 1c).

Now, to manipulate the life history of the consumer, we added the parameter *ε* to the consumer’s equation. The consumer is fast when *ε* is one and slow when *ε* is tiny.

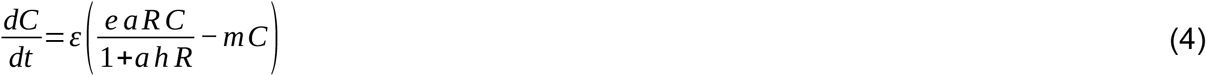

Because *ε* scales the consumer’s intrinsic growth rate (*e·a*_*max*_ *– m* where *a*_*max*_ is 1/*h*, see S.I. Section “Scaling of the consumer’s growth rate by ε”) but preserves the isocline geometry of the C-R model, we can manipulate the life history of the consumer along a slow-fast continuum (relative to the resource) (Figure 1a) while controlling the underlying deterministic skeleton.

For all analyses, *r =* 2.0, *k =* 3.0, *a* = 1.1, *h* = 0.8, and *m* = 0.4.

### Stability along the slow-fast continuum

We used the coefficient of variation to measure stability as the consumer’s life history was varied from fast to slow (1/ε was varied from 1.0 to 1000). We used a flow-kick approach to add stochasticity (versus a stochastic ordinary differential equation approach) [32]. In this flow-kick approach, we kick the consumer variable every unit of time and let the model flow (integrate) without noise until the next kick. The kick is generated from normally distributed noise (with mean 0.0 and standard deviation 0.01). For each value of 1/ε, we ran 50 simulations of 24,000 time units (package DifferentialEquations.jl v6.20.0, Algorithms: Vern7 & Rodas4 with automatic stiffness detection). From these simulations for each value of 1/ε, we calculated the average coefficient of variation (CV) after removing the first 2000 time units. We did this for the efficiency values (*e*) of 0.5 (non-excitable deterministic dynamics with real eigenvalues) and 0.71 (excitable deterministic dynamics with complex eigenvalues). Note, any simulations that landed on the axial solution (where the resource isocline intersects the Consumer = 0 axis) were removed from the average coefficient of variation calculation.

To unpack the CV result as the consumer’s life history is slowed, we examined the dynamical behaviours that drove the change in CV. Three general dynamical behaviors occur depending on whether the C-R interaction is excitable (complex eigenvalues) or non-excitable (completely real eigenvalues) (see Figure 2a). The first dynamical behaviour is quasi-cycling and occurs only when the C-R interaction is excitable (see Box 1 and Figure 2b). Quasi-cycling is when stochastic perturbations resonate with the excitability of the C-R model to extend the range of cyclic dynamics (see Box 1 and [21,22]). Compared to the deterministic limit cycles, these quasi-cycles generally do not threaten persistence because quasi-cycles exhibit smaller variation. Nevertheless, because slowing the consumer’s life history reduces the excitability of the C-R interaction (see S.I. Section “Slowing the consumer decreases excitability”), we may see an impact of slowing the consumer’s life history on the stability (CV) of the C-R interaction. The second dynamical behavior is quasi-canards which can happen for both excitable and non-excitable C-R interactions (see Box 1 and Figure 2b). A quasi-canard is similar to a deterministic canard. A deterministic canard occurs after the Hopf bifurcation (i.e. to the left of the resource isocline maximum) and does not require stochastic noise. In contrast, a quasi-canard occurs before the Hopf bifurcation (i.e. to the right of the resource isocline maximum) and requires stochastic noise (see Box 1). Because quasi-canard trajectories are quite large, stability will decrease markedly (i.e. CV will increase). The third dynamical behaviour is what we term stretched wandering and occurs for non-excitable C-R interactions only (see Figure 2b). The dynamics are stretched along the resource isocline with no obvious cycling.

**Figure 2.**
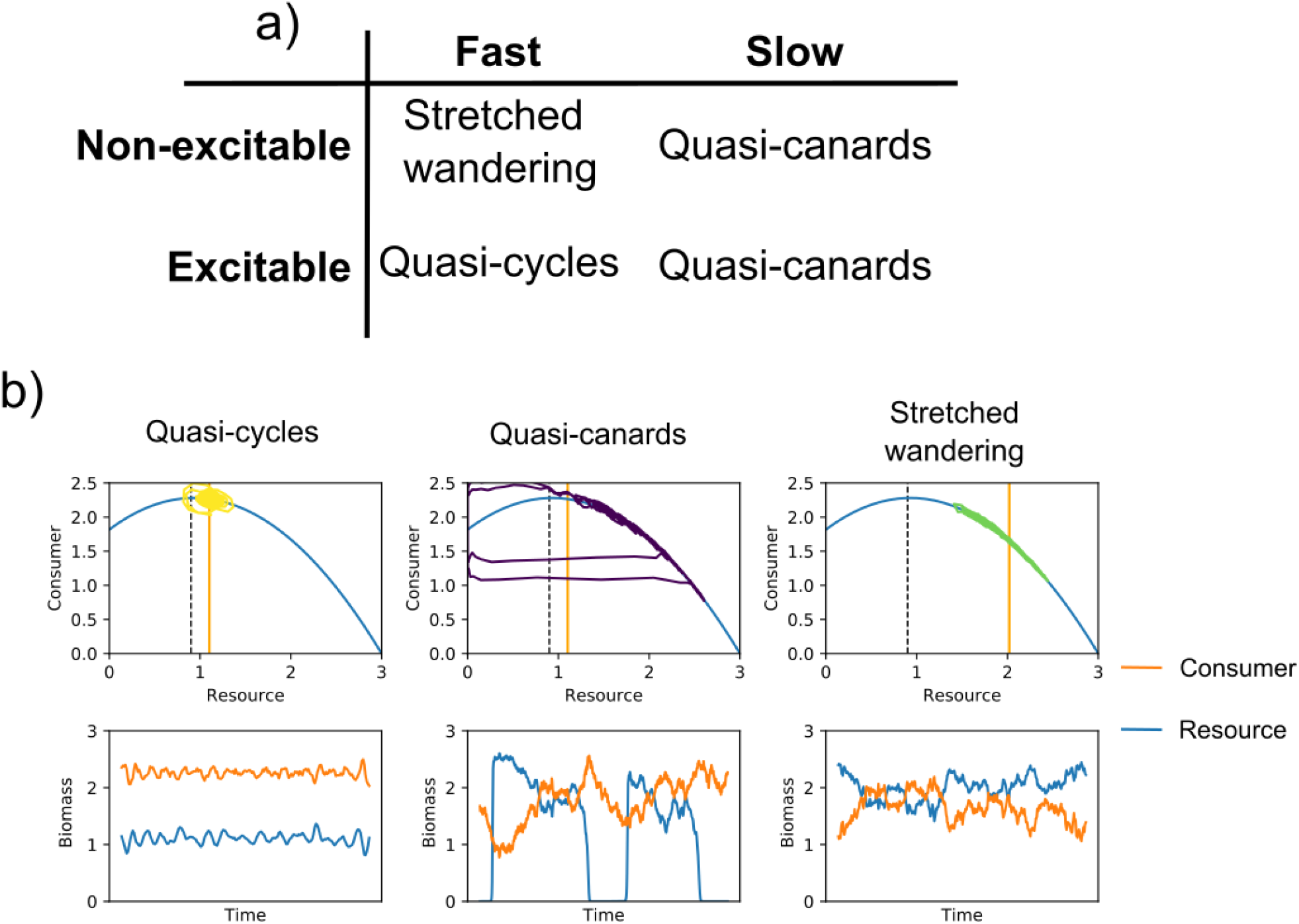
a) Table of dynamical behaviours when the C-R interaction is non-excitable and excitable and when the consumer’s life history is fast or slow. b) Phase diagrams and time series of quasi-cycles (yellow), quasi-canards (purple), and stretched wandering (green). Dashed vertical lines in the phase diagrams denote where the deterministic Hopf bifurcation occurs. Note, the standard deviation of the noise was increased to 0.05 to help emphasize what quasi-cycles, quasi-canards, and stretched wandering look like.

### Quasi-cycles

To examine how slowing the consumer’s life history impacted quasi-cycles, we ran 100 simulations each of the C-R model with 1/ε values of 1 and 10 and where, again using the flow-kick approach, the consumer variable was perturbed every time unit with normally distributed noise (with mean 0.0 and standard deviation 0.01) (package DifferentialEquations.jl v6.20.0, Algorithms: Vern7 & Rodas4 with automatic stiffness detection). To examine the initial decrease in CV as the consumer’s life history was slowed, we chose 1/ε values of 1 and 10. We calculated the autocorrelation of the last 1000 time units of each simulation for lags between 0 and 40 (function autocor, package StatsBase v0.33.13). We then calculated the average autocorrelation value for each lag across the 100 simulations for each simulation set with a different 1/ε value. We did this for the efficiency values (*e*) of 0.5 (non-excitable) and 0.71 (excitable). As suggested by Pineda-Krch et al. [21], we used autocorrelation to identify the occurrence of quasi-cycles. An autocorrelation function (ACF) from quasi-cycles would show pronounced amplitude oscillations that decrease in amplitude with increasing lags. No oscillations in the ACF indicate that quasi-cycles are not occurring and constant ACF oscillations indicate noisy period cycling. An ACF with smaller damped oscillations indicates that the periodicity of the quasi-cycles is being removed. Note, that in our ACF figures, the mean of each time series was removed from each time point value. In the ACF figures in Pineda-Krch et al. [21], the mean of each time series was not removed from each time point value. Thus, our figures look different (our ACF scale goes from -1 to 1 whereas Pineda-Krch et al.’s [21] scale goes from 0.95 to 1). Our method makes spotting the damped oscillations easier.

### Quasi-canards

To examine how slowing the consumer’s life history impacted quasi-canards, we examined the prevalence of quasi-canards in stochastically perturbed simulations of the C-R model with varying ε and efficiency values. First, we examined the prevalence of quasi-canards under white noise stochasticity. We ran 1000 simulations with 24,000 time units for each value of 1/ε varied from 6.667 to 1000, where, using the flow-kick approach, the consumer variable was perturbed every time unit with normally distributed noise (with mean 0.0 and standard deviation 0.01) (package DifferentialEquations.jl v6.20.0, Algorithms: Vern7 & Rodas4 with automatic stiffness detection). Our lower bound for 1/ε was set at 6.667 because no quasi-canards can be found below this point. This was done for efficiency values of 0.5 (non-excitable) and 0.71 (excitable). For each combination of *ε* and efficiency value, we calculated the proportion of simulations that exhibited at least one quasi-canard. To calculate the proportion of simulations with quasi-canards, we created an algorithm to check whether a time series contained a quasi-canard. More details of this algorithm can be found in the Supporting Information (S.I. section “Explanation of quasi-canard finder algorithm”), but in short the algorithm includes a return map at the maximum point of the resource isocline which canards and quasi-canards must pass through. The algorithm also includes boxes along the attracting and repelling manifolds (the right side of the resource isocline and the consumer axis respectively) through which a quasi-canard should pass.

Second, we examined how reddened noise impacts the prevalence of quasi-canards. For two values of 1/ε (12.66 and 250.0) and for two values of efficiency (0.5 and 0.71), we varied the stochasticity from white to red noise and measured the proportion of simulations that exhibited at least one quasi-canards. Note, we also measured the proportion of simulations that exhibited no quasi-canards but landed on the axial solution (where the resource isocline intersects the Consumer = 0 axis) because landing on the axial solution readily occurs under red noise. We ran 1000 simulations with 6,000 time units for each value of 1/ε, efficiency, and noise autocorrelation (package DifferentialEquations.jl v6.20.0, Algorithms: Vern7 & Rodas4 with automatic stiffness detection). We used an autoregressive model of order 1 (AR_1_) to create reddened noise with autocorrelation varying from 0 (white noise) to 1 (red noise). We scaled the variance of the red noise to the variance from the original white noise using the technique in Wichmann et al. [33] where the ratio of white noise to red noise variances scales individual noise values in the red noise sequence.

All analyses were done using julia version 1.7.0. [34]

## Results

### Stability along the slow-fast continuum

When the C-R model is non-excitable (e = 0.5), slowing the consumer’s life history increases the coefficient of variation (Fig. 3). When the C-R is highly excitable (e = 0.71), slowing the consumer’s life history initially decreases then increases the coefficient of variation (Fig. 3).

**Figure 3.**
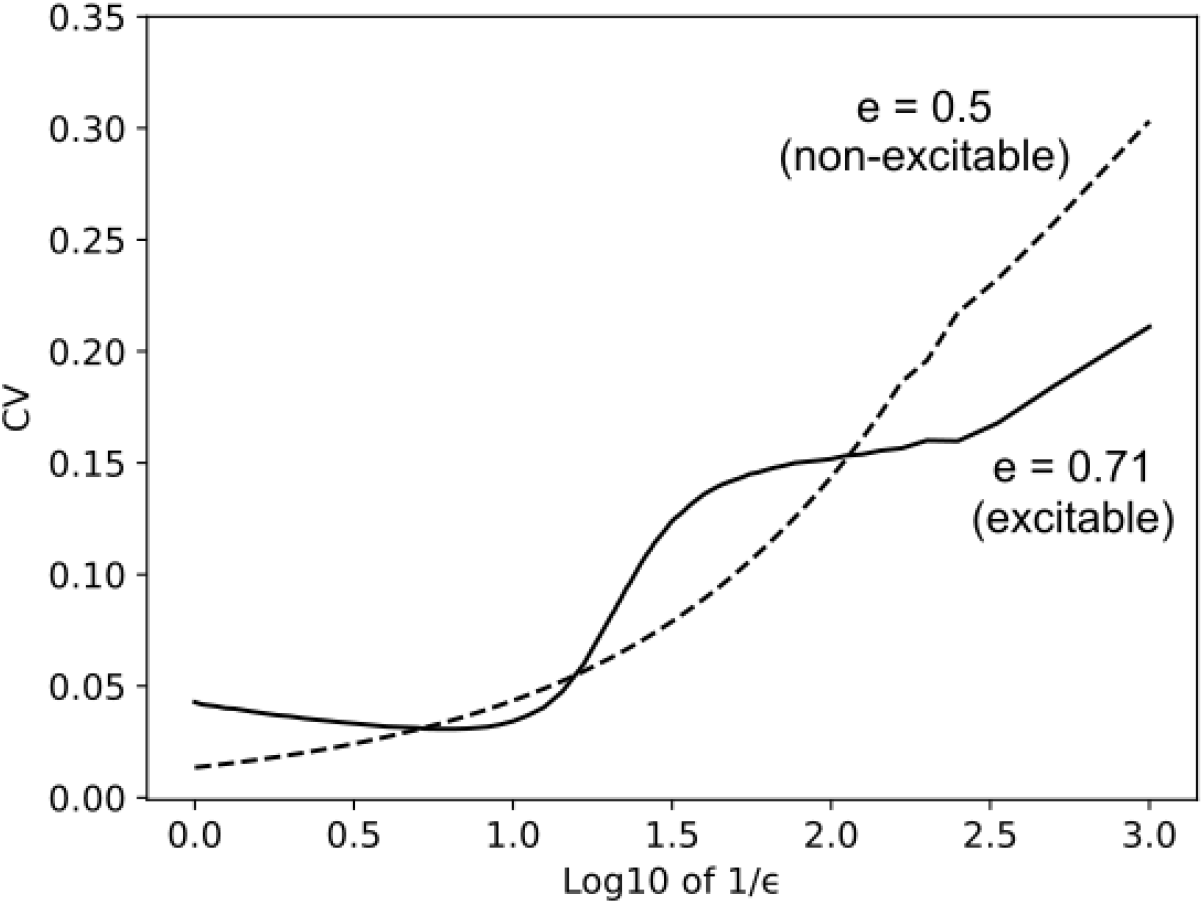
Average coefficient of variation for time series of Rosenzweig-MacArthur C-R model with efficiency of either 0.5 (non-excitable) or 0.71 (excitable) along a continuum of 1/ε from 1 to 1000 (50 simulations per value of 1/ε and efficiency). Note, reducing life history speed is first modestly stabilizing, in effect removing the oscillatory potentials of the underlying excitable model before life history slowing ultimate generates large increases in CV that likely produce population disappearances (R∼0 during trajectories) due to quasi-canards.

### Quasi-cycles

When the C-R model is non-excitable (e = 0.5), quasi-cycles are never found regardless of how slow the consumer’s life history is (Fig. 4 a, c, & e). In contrast, when the C-R model is highly excitable (e = 0.71) quasi-cycles are found when 1/*ε* = 1.0 (Fig. 4b & d). With a slowing of the consumer’s life history (now 1/*ε* = 10) the average autocorrelation function (ACF) line flattens out indicating reduced manifestation of quasi-cycles (Fig. 4b & f).

**Figure 4.**
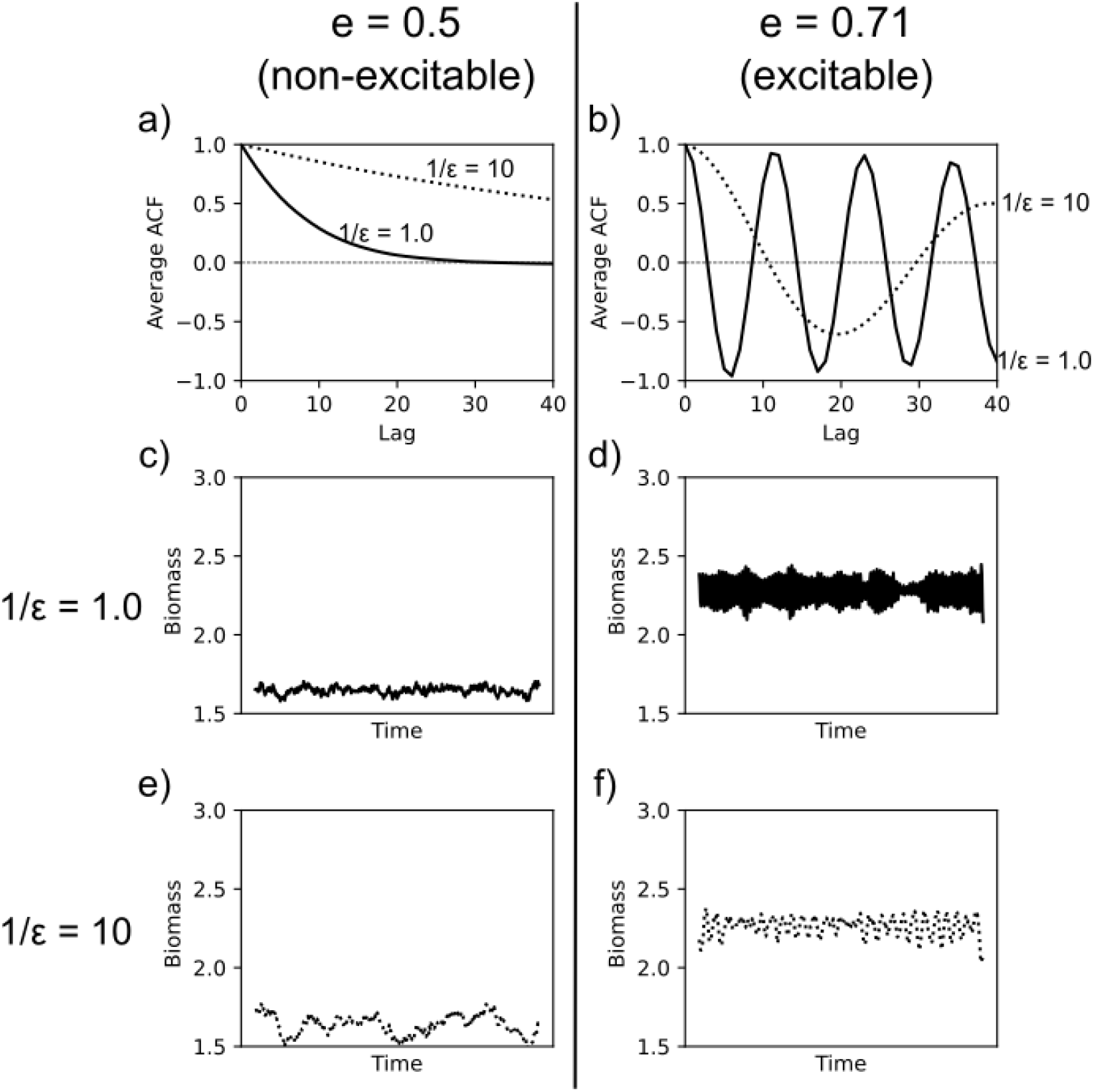
a) & b) Average ACF for each lag value for 100 simulations of the C-R model perturbed each time unit by normally distributed noise (with mean 0.0 and standard deviation 0.01) with 1/ε values of 1 and 10 and with efficiency values of 0.5 (non-excitable) and 0.71 (excitable) respectively. c) & d) Time series of the consumer with 1/ε value of 1 and efficiency values of 0.5 and 0.71 respectively. e) & f). Time series of the consumer with 1/ε value of 10 and efficiency values of 0.5 and 0.71 respectively.

### Quasi-canards

When the C-R model is non-excitable (e = 0.5), quasi-canards can be found under white noise but require a very slow consumer life history (Fig. 5a). When the C-R model is highly excitable (e = 0.71), quasi-canards can be easily found along the gradient of fast to slow consumer life histories (Fig. 5b). Reddening the noise increases the likelihood of finding both quasi-canards and the axial solution regardless of the consumer’s life history (Fig. 5 c, d, e, f). Note for a highly excitable C-R interaction, landing on the axial solution is less likely than for a non-excitable C-R interaction because there is more phase space that the dynamics must traverse through. Slowing the consumer’s life history down increases the likelihood of the highly excitable C-R interaction landing on the axial solution.

**Figure 5.**
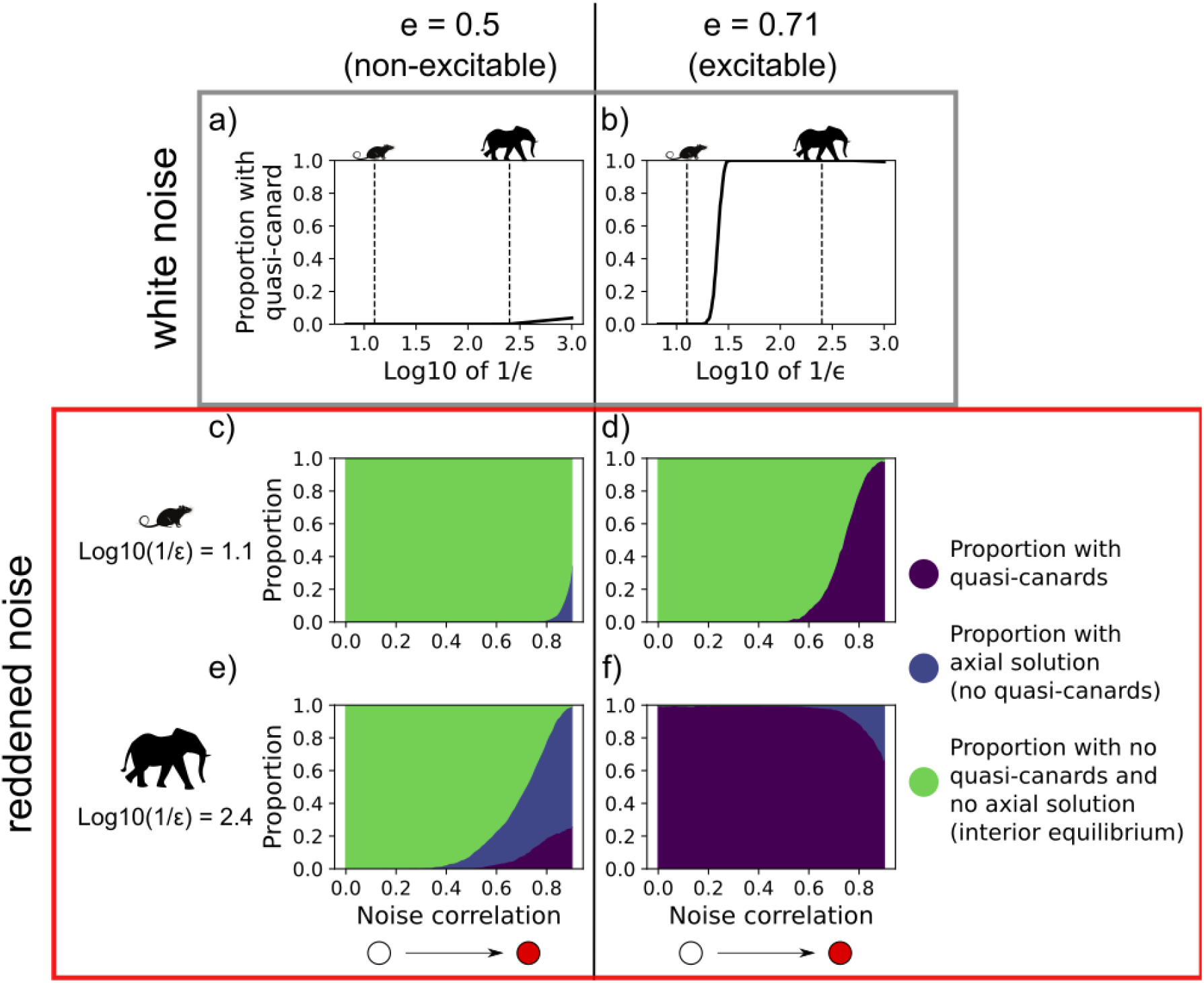
a) & b) Proportion of 1000 simulations under white noise per value of 1/ε that exhibited quasi-canards with constant efficiency values of 0.5 (non-excitable) & 0.71 (excitable) respectively. Vertical dashed lines correspond to log10(1/ε) values used in c), d), e), & f). c), d) Proportion of 1000 simulations per value of log10(1/ε) = 1.1 that exhibited quasi-canards or axial solution or neither with constant efficiency values of 0.5 & 0.71 respectively and with noise correlation (AR1 process) varied from 0.0 to 0.9. e), & f) Proportion of 1000 simulations per value of log10(1/ε) = 2.4 that exhibited quasi-canards or axial solution or neither with constant efficiency values of 0.5 & 0.71 respectively and with noise correlation (AR_1_ process) varied from 0.0 to 0.9.

## Discussion

Using the simple technique of changing the time scales within the Rosenzweig-MacArthur consumer-resource (C-R) model, we set up a biologically motivated mathematical experiment exploring the stability of faster and slower life histories under white to red stochastic noise. Here, we experimentally manipulated the consumer’s intrinsic growth rate while maintaining the underlying qualitative dynamical conditions. When the energy flux is high, we find that initially slowing the consumer’s life history increased stability through reduced manifestation of quasi-cycles (alternatively, when energy flux is low, slowing the consumer’s life history always decreased stability). However, further slowing the consumer’s life history decreased stability through increasing the likelihood of quasi-canards and population disappearances. Furthermore, we found that increasing the autocorrelation of noise tended to increase the likelihood of quasi-canards and population disappearances. Finally, Yodzis & Innes’s [23] biologically plausible C-R model similarly exhibits quasi-canards (see S.I. Section “Biologically plausible parameters” & S.I. Figure 5). Consequently, our results are general and suggest that such instability ought to occur under increasingly reddened perturbations arising from climate change.

Our results illuminate a gradient in stability along the slow-fast life history continuum that is dependent on energy flux. If the C-R interaction is highly excitable, initial slowing of the consumer’s life history can increase stability. Stability increases because slowing the consumer reduces the excitability of the C-R interaction which reduces the manifestation of the quasi-cycles (see S.I. Section “Slowing the consumer decreases excitability”). If the C-R is non-excitable or moderately excitable, initial slowing of the consumer’s life history decreases stability (although slowing the consumer’s life history does reduce quasi-cycles for the moderately excitable C-R interaction). Here, perturbations push a slow growth consumer away from the equilibrium and thus the coefficient of variation continually increases. For all types of C-R interaction, when the consumer’s life history is sufficiently slow, additional perturbations multiply the effect of each perturbation resulting in a consumer biomass that is “far from equilibrium” and prone to the distant nonlinear effects of the C-R dynamics (i.e, the quasi-canard). Overall, there appears to be a gradient in stability along the slow-fast life history continuum that is mediated by energy flux.

The combination of life history and red noise significantly increased the opportunities for non-local nonlinear dynamics to be expressed. Reddening the noise in our C-R model increased the likelihood of quasi-canards (and landing on the axial solution) for both fast and slow organisms. However, the onset of quasi-canards occurred with less autocorrelation for faster organisms. To understand this pattern, we must examine the relative time scales of the autocorrelation and the system population processes [26]. Slowing the life history is effectively increasing the time scale of the population response processes, and thus the system will stay for longer in the phase space region the system was pushed into after a perturbation. Increasing the autocorrelation increases the time scale of the perturbations, and thus the perturbations effectively are magnified over time.

To unpack the idea of the magnification of the perturbations, we can use similar research on how multiple discrete disturbances can kick dynamics out of basins of attraction to produce different dynamical outcomes (flow-kick dynamics: Meyer et al. [32]). In the flow-kick framework, models are kicked at discrete time points and are allowed to flow (integrate) without kicks in between the discrete time points. Meyer et al. [32] found that rare but large disturbances can have the same effect as frequent and small disturbances. Reddened noise is technically lots of frequent and small disturbances (the kicks in flow-kick dynamics). But because of the autocorrelation pushing dynamics far from their attracting equilibria, reddened noise has the same effect as a large disturbance. Furthermore, slowing life histories reduces the relative time available for organisms to respond to the disturbances (the flow in flow-kick dynamics), thus pushing dynamics far from their attracting equilibria.

Overall, we can use the analogy of a rusty ratchet to understand how slow life histories and reddened noise interact. White noise is akin to a ratchet that can spin in any direction (without a pawl), and red noise is akin to a ratchet with a pawl that can spin in only one direction for a period of time (Figure 6). Because the reddened noise has a tendency to produce similar values for a period of time, reddened noise consistently pushes the dynamics of the C-R model far from the local area around the stable equilibria. Whereas the fast life history is akin to oil in the ratchet, the slow life history is akin to rust in the ratchet which slows the spinning speed and in the system increases the time required for trajectories to return to the local area around the equilibria. To describe this process, we use the term noise-induced ratchet effects inspired by the rate-induced critical transition literature [35,36]. In a rate-induced critical transition, canards can be produced when the equilibrium shifts at a slow rate. The dynamics enter non-local non-linear dynamics because the equilibrium has been pulled from under the dynamics. For the quasi-canard, the dynamics enter non-local non-linear dynamics because the dynamics have been pushed by the noise (thus quasi-canards are noise-induced critical transitions). When slow life histories are combined with red noise, we get noise-induced ratchet effects.

**Figure 6.**
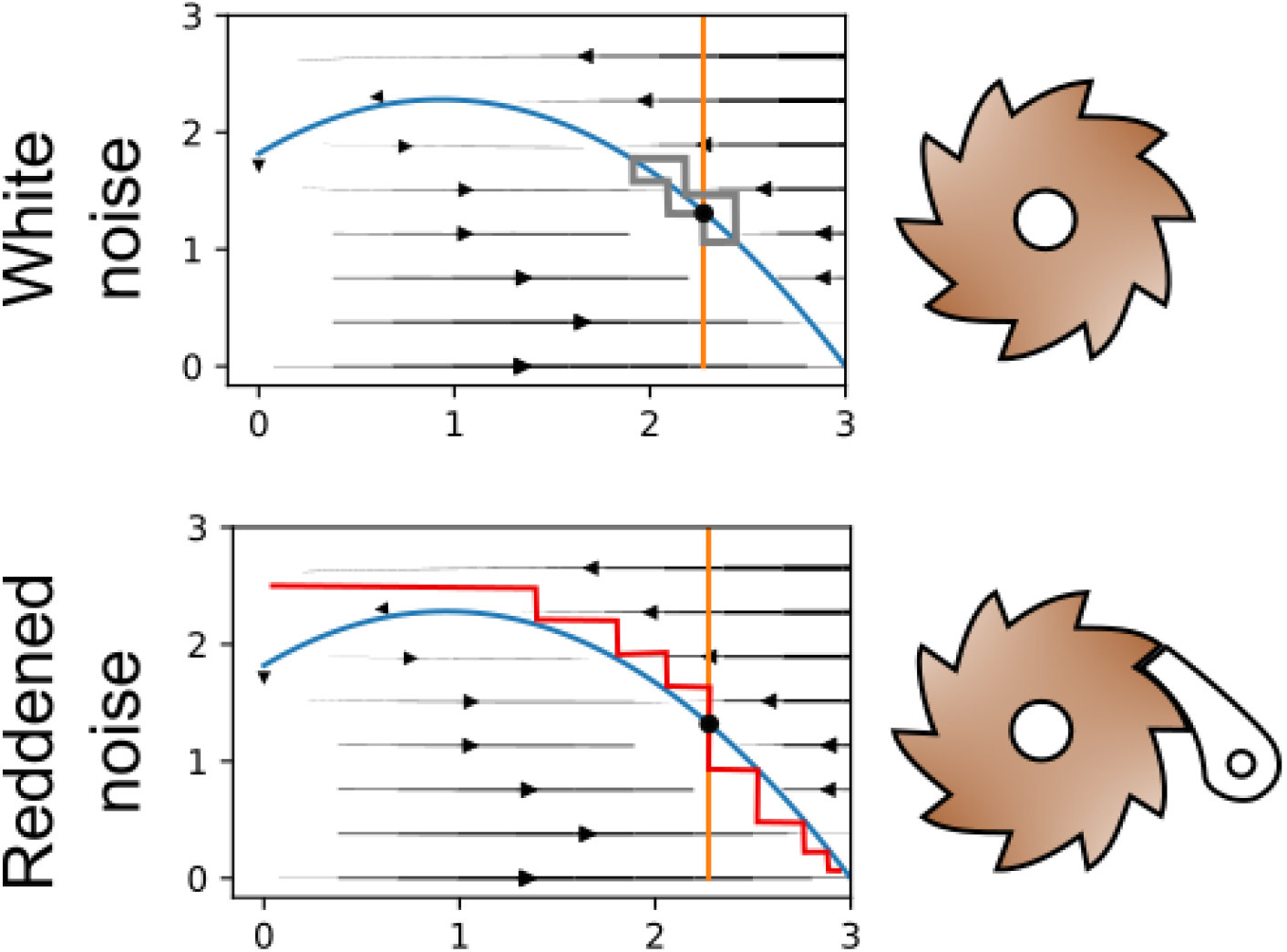
Illustration of simplified trajectories with white noise (top) and red noise (bottom) together with the consumer and resource isoclines and vector field when the consumer’s life history is slow (1/*ε* is large). We use the analogy of a rusty ratchet to illustrate the interaction of slow life histories with reddened noise. White noise is similar to a ratchet wheel without the pawl (can spin in any direction) and reddened noise is similar to a ratchet with the pawl (can spin in only one direction for a period of time). Slow life history is akin to rust in the ratchet which slows the spinning speed.

Intriguingly, our examination of slow life histories with stochasticity exhibiting sudden population disappearances is a further example of how stochasticity is immensely useful in ecological research to understand the full nonlinear dynamics of ecosystems [20,26,29]. First, stochasticity can act to uncover underlying processes (Boettiger [26] coined the phrase *noise the informer* for this phenomenon). Similar to quasi-cycles where stochastic resonance is visible in advance of a Hopf bifurcation, the quasi-canards also occur in advance of the Hopf bifurcation after which deterministic canards occur. Although noise has the same effect of uncovering an imminent Hopf bifurcation leading to either deterministic cycles or canards, the mechanism producing the quasi-cycles and quasi-canards are different. The quasi-cycles are stochasticity interacting with local nonlinear dynamics (excitability). The quasi-canards are slow life histories plus reddened noise pushing dynamics towards non-local nonlinear dynamics.

Second, stochasticity can be used to reveal differences in stability from what our normal linear stability analysis would find and indeed linear stability analysis would not have predicted the highly destabilizing behaviour for slow consumers. One method to reveal these differences in stability is through using the mathematical tools of the potential function or quasi-potential where potential functions and quasi-potentials can be simplified conceptually to the ball and cup analogy [37]. In this analogy, the state of a system is the position of the ball rolling around a surface (the cup) with minimums being attractors. The potential and quasi-potential are the cup. The potential function can be found for any system that exhibits solely fixed point attractors (gradient systems). Quasi-potentials can be found for gradient and non-gradient systems (the C-R model is a non-gradient system because it exhibits cycling). The quasi-potentials for our model show the stretching of the quasi-potential along the resource isocline due to stochasticity with low efficiency and a fast consumer life history, then the quasi-canard shape when efficiency is large enough, and finally the flattening of the quasi-potential quasi-canard shape with a slow consumer life history (see S.I. Figure 6). Linear stability analysis would solely focus on the tiny region around the intersection of the consumer and resource isoclines. In contrast, the flattened quasi-potentials reveal other possible dynamics (quasi-canards) in addition to the stable interior equilibria. Overall, stochasticity can both produce and reveal dynamical outcomes not predicted by normal linear stability analysis.

## Conclusion

By varying growth rates along the slow-fast life history continuum within the classic Rosenzweig-MacArthur model, we have shown that life history has interesting stability consequences for the system. Slowing the life history can initially increase stability up to a point in the face of many tiny perturbations, but further slowing can dramatically decrease stability and increase the potential for highly variable dynamics and population disappearances. Noise-induced ratchet effects occur when positively correlated noise is added to an already slowed consumer. This noise-induced ratchet effect is another mechanism that selects against slow life histories leading to greater proportions of species with fast life histories. Our study examines a single C-R interaction, which is a fundamental component of food web theory. Moving forward, the interaction of many more organisms with varying life histories is required for a more comprehensive understanding of life history and stability. Furthermore, other examples of non-local nonlinear dynamics should be explored, especially in the context of slow life histories and reddened noise. Taken together, we have shown how life history along the slow-fast life history continuum can impact the stability of systems and shown how human-caused reddened noise will disproportionately impact slow living organisms through noise-induced ratchet effects and population disappearances.

## Acknowledgments

We wish to thank members of the McCann laboratory for thoughts and feedback.

## Funding

CJGG was supported by a Natural Sciences and Engineering Research Council of Canada (NSERC) CGS-D and a Ontario Graduate Scholarship. Financial support was provided by an NSERC Discovery Grant (400353) and a CFREF Food from Thought grant (499075) to K. S. McCann.

## Author Contributions

All authors contributed to the idea generation and analysis of the model. CJGG wrote the first draft and all authors contributed to editing the manuscript.

## Code accessibility

All code to reproduce the above analyses and figures are publicly available on [GitHub](https://github.com/cgreysongaito/SlowFast_SuddenPopulationDisappearances) and have been archived on [Zenodo](https://zenodo.org/record/7246515) (version 2.0).

## Supporting Information

### Scaling of the consumer’s growth rate by *ε*

To see the impact of ε on the consumer’s growth rate let’s rearrange the parameters in the model to get the functional response into a monod equation

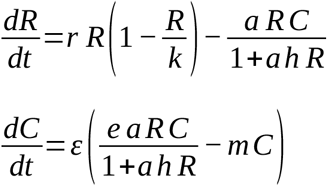

becomes

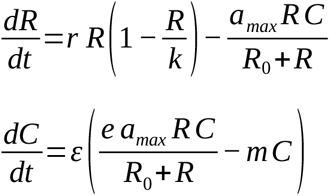

where 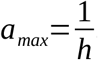 and 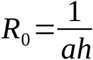.

From this version of the model, we can see that ε scales *e·a*_*max*_ *– m* (the consumer’s growth rate).

### Isoclines of Rosenzweig-MacArthur consumer-resource model

**Figure S.I.1.**
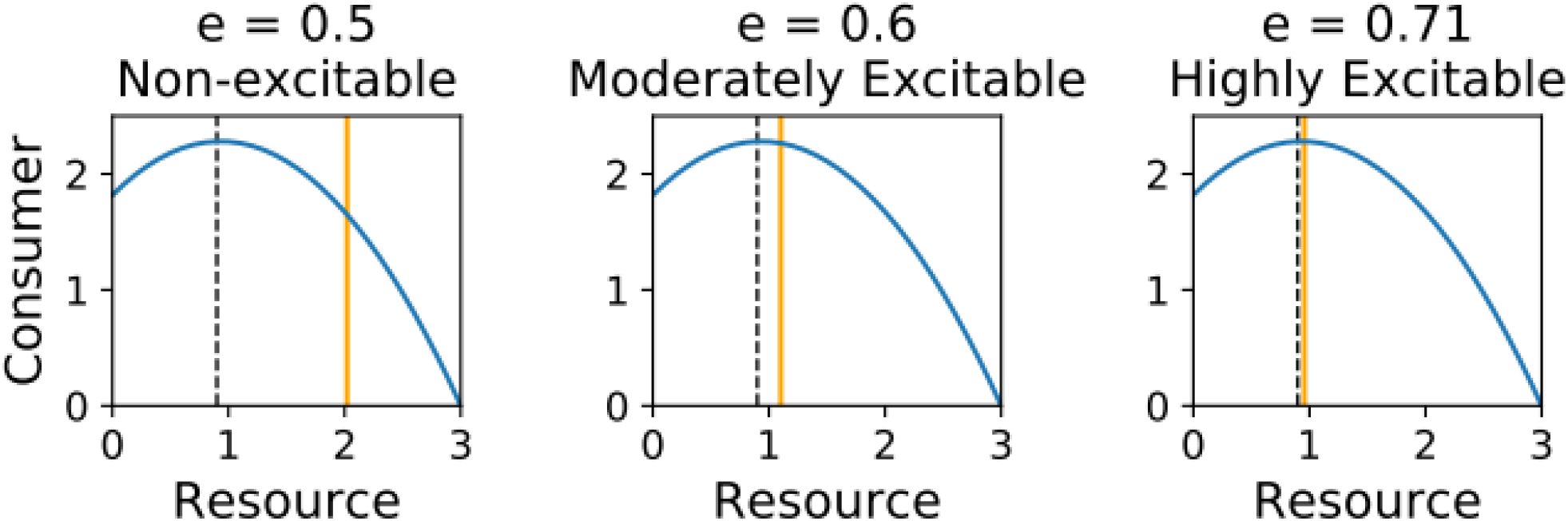
a), b), c) resource and consumer isoclines with efficiency values of 0.5, 0.6, 0.71 respectively. Dashed lines depicts where the Hopf birfurcation occurs

### Moderately excitable consumer-resource interaction

**Figure S.I.2.**
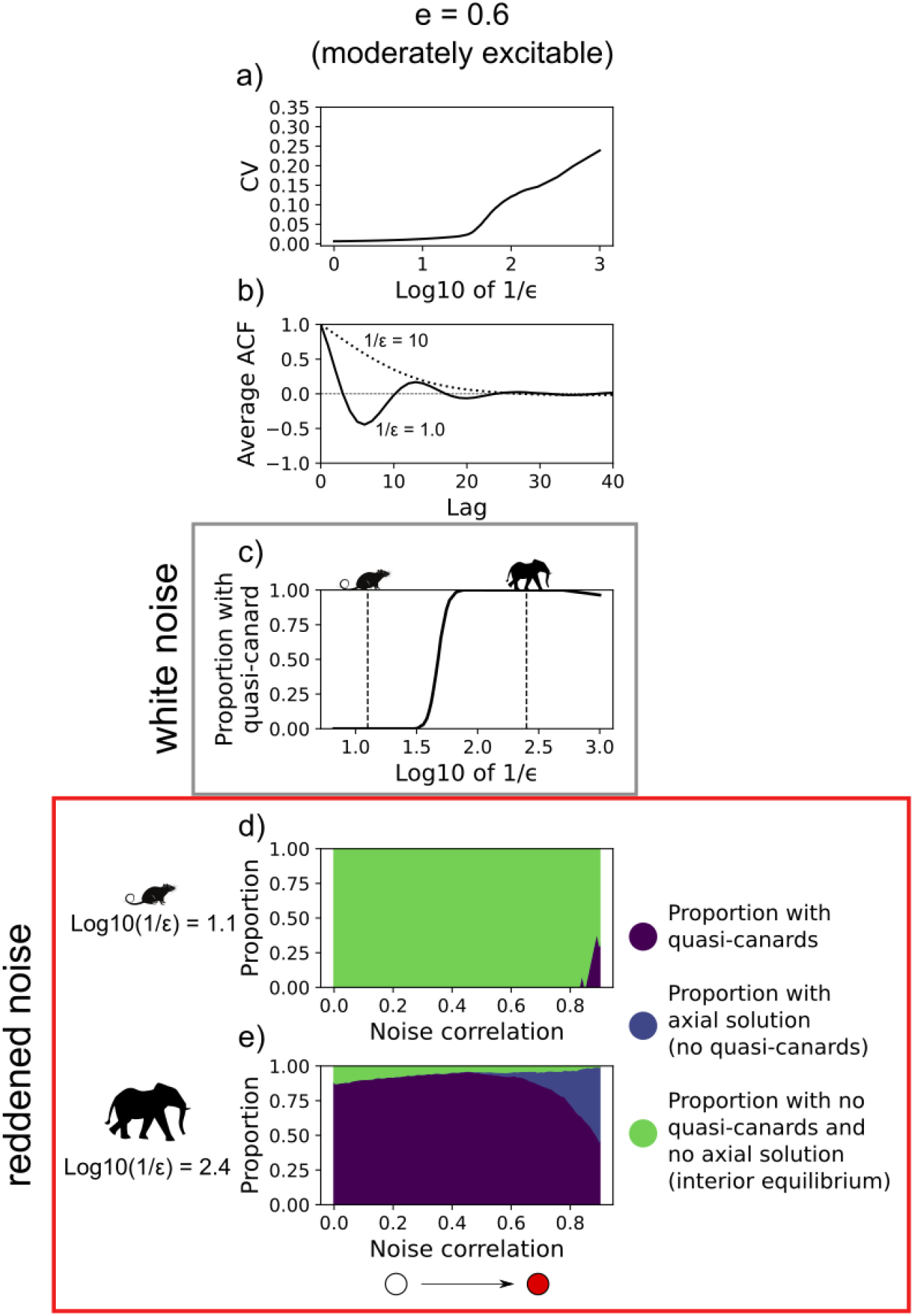
a) Average coefficient of variation for time series of Rosenzweig-MacArthur C-R resource model with efficiency of 0.6 (moderately excitable) along a continuum of 1/ε from 1 to 1000 (50 simulations per value of 1/ε and efficiency). b) Average ACF for each lag value for 100 simulations of the C-R model perturbed each time unit by normally distributed noise (with mean 0.0 and standard deviation 0.01) with 1/ε values of 1 and 10 and with an efficiency value of 0.6 (moderately excitable). c) Proportion of 1000 simulations under white noise per value of 1/ε that exhibited quasi-canards with an efficiency value of 0.6 (moderately excitable). Vertical dashed lines correspond to 1/ε values used in d), & e). d), e) Proportion of 1000 simulations per value of log10(1/ε) = 1.1 and log10(1/ε) = 2.4 respectively that exhibited quasi-canards or axial solution or neither with an efficiency value of 0.6 and with noise correlation (AR1 process) varied from 0.0 to 0.9.

### Slowing the consumer decreases excitability

Below, we numerically and analytically prove that slowing the consumer’s life history (increasing 1/*ε*) decreases the excitability of the C-R interaction. By reducing the excitability, we mean that the divide between real and complex eigenvalues (from the linear stability analysis of the interior equilibrium) moves towards the Hopf bifurcation.

#### Numerical proof

We numerically calculated the real to complex divide for each value of 1/ε between 0.01 and 10 — with a step size of 0.001 in ε (package LinearAlgebra.jl). For each value of 1/ε, we found the efficiency value, *e*_*R/C*_, where the real/complex divide occurs (i.e. after 1/ε is set, the efficiency value is increased from the efficiency value that produces the transcritical bifurcation until the eigenvalues switch from real to complex, akin to finding the value at which the tip of the checkmark occurs in Gellner and McCann [12]). This efficiency value was then used to calculate the proportion of efficiency “parameter space” that produces real eigenvalues (Proportion Real) by subtracting the real/complex divide efficiency value (*e*_*R/C*_) from the efficiency value at the Hopf bifurcation (*e*_*Hopf*_) and then dividing this value by the efficiency parameter distance between the deterministic Hopf (*e*_*Hopf*_) and transcritical (*e*_*transcritical*_) bifurcation efficiency values (note *ε* does not change where the Hopf and transcritical bifurcation occur):

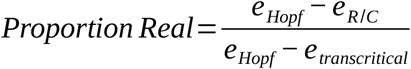

Increases in 1/*ε* move the real-complex divide towards the Hopf bifurcation and thus, increase the proportion of efficiency “parameter space” that produces real eigenvalues (Figure SI1). In other words, increasing 1/*ε*, reduces the excitability of the system.

**Figure S.I.3.**
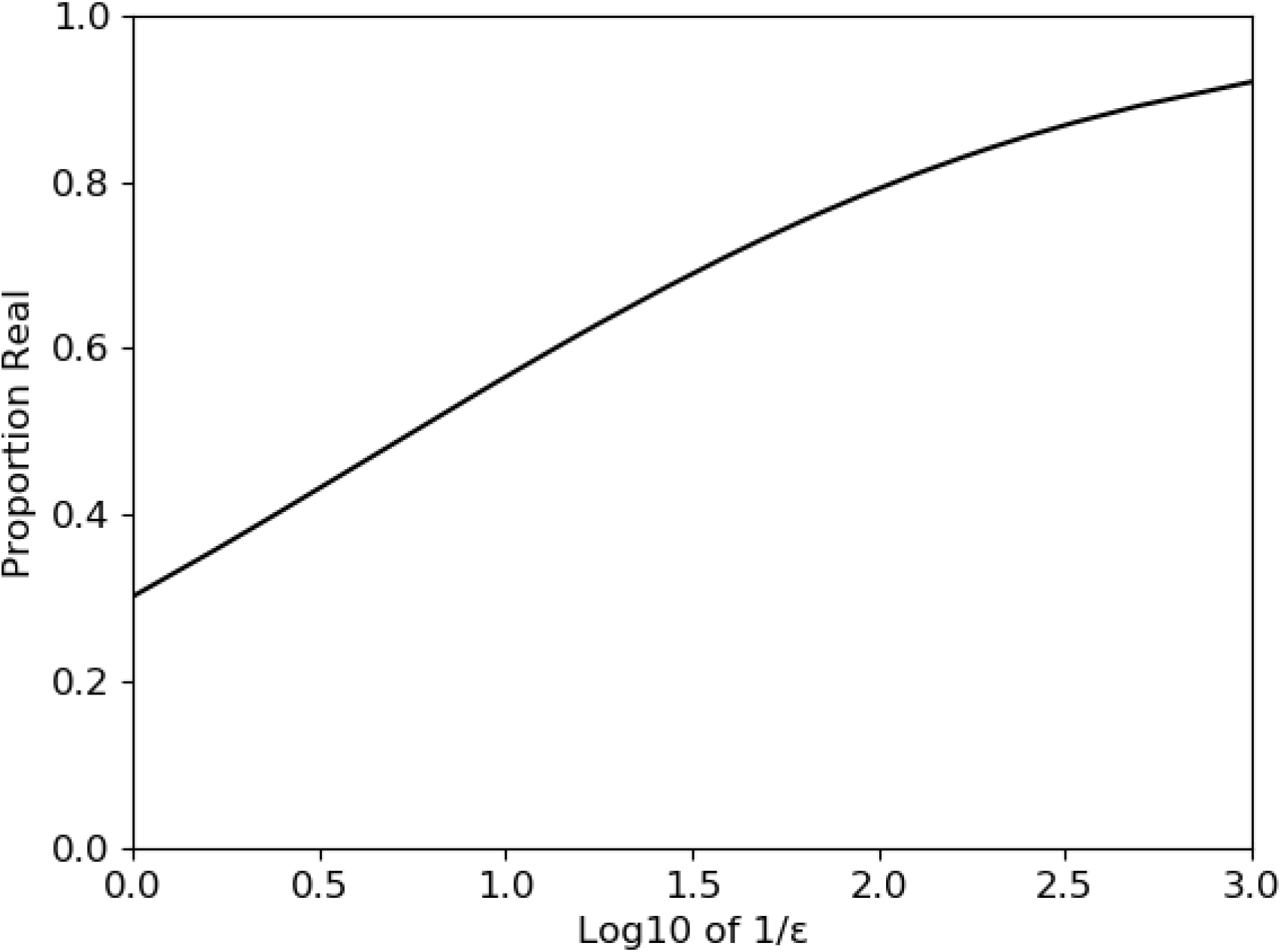
Proportion of efficiency “parameter space” that produces completely real eigenvalues for each 1/ε value where the full efficiency “parameter space” corresponds to the distance between the efficiency values that produce the transcritical and Hopf bifurcation.

#### Analytical Proof

We also proved, using the non-dimensional type I version of the Rosenzweig-MacArthur C-R model, that increasing 1/*ε*, increases the efficiency value where the real/complex divide occurs and thus decreases excitability.

With change of variables

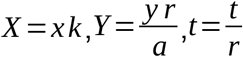

and with non-dimensional parameters 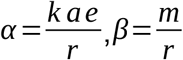 the non-dimensionalized form of the type I functional response C-R model is:

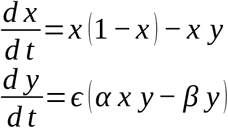

Equilibria exist at

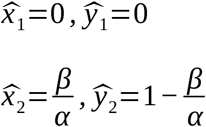

The jacobian of the model is

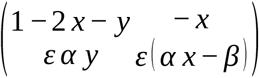

Inputting the interior equilibrium, 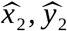, into the jacobian returns

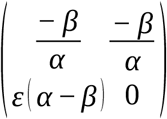

Using the trace and determinant of this jacobian matrix we can get the characteristic polynomial and the quadratic equation to solve for the eigenvalues:

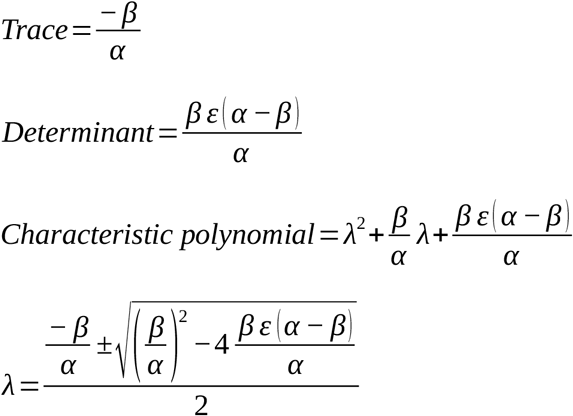

We are determining the boundary of real to complex eigenvalues, thus we must examine what is inside the square root of the quadratic equation:

When 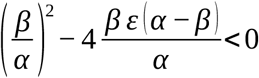 the eigenvalues are complex

We can solve for α to find what parameter values produce α at the real/complex divide

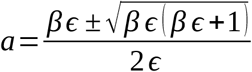

We can ignore the minus square root part (because 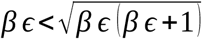 always and we get a negative alpha value which is impossible biologically).

Thus, we concentrate on

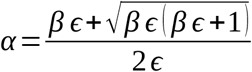

We can differentiate the above equation with respect to *ε* to find out how the α value (at which the real/complex divide occurs) changes.

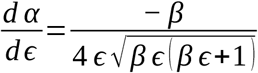

which is always negative when *β* and *ε* are positive (biologically they have to be).

Therefore, if we decrease *ε* (slowing the consumer by increasing 1/*ε*), the *α* value — at which the real/complex divide occurs — increases. Converting *α* back into its original dimensional parameters, we see that if *k* and *a* are kept constant, *e* must increase to increase the non-dimensional *α* parameter.

### Explanation of quasi-canard finder algorithm

The algorithm checks that the trajectory has the characteristics of a quasi-canard. Thus, the algorithm includes a return map at the maximum point of the resource isocline where canards and quasi-canards must pass through. The algorithm also includes boxes along the attracting and repelling manifolds (the right side of the resource isocline and the consumer axis respectively) through which a quasi-canard should pass. The quasi-canard passes through these checks in a particular order and so the algorithm ensures the order is correct. Below are the six steps that the quasi-canard finder algorithm goes through. The full code can be found in slowfast_canardfinder.jl of the Github repository.

1. The algorithm finds all the points in the time series where the next sequential point creates a vector that intersects with a line that sits at the Hopf bifurcation point on the Resource isocline (the maximum of the Resource isocline). The line has a length of 5% of the Hopf bifurcation point above and below the Hopf bifurcation point. If no points are found, the algorithm does step six. If points are found, the points are collated and passed to the next step.
2. The algorithm then takes all of these points and moves along the time series after these points to identify the first point within a box that sits between the Hopf bifurcation point and where the Resource isocline intersects with the Consumer axis. The box has a width of 0.1. If no points are found, the algorithm does step six. If points are found, the points are collated and passed to the next step.
3. The algorithm then takes all of these points and moves along the time series after these points to identify the first point within a box that sits between the 0 consumers and 80% of where the Resource isocline intersects with the Consumer axis. The box has a width of 0.1. If no points are found, the algorithm does step six. If points are found, the points are collated and passed to the next step.
4. The algorithm then takes all of these points and moves along the time series after these points to identify the first point that sits close to the resource isocline. If no points are found, the algorithm does step six. If points are found, the points are collated and passed to the next step.
5. The algorithm then takes all of these points and repeats step 1 to ensure a full cycle of the quasi-canard. If the return map check is passed for the second time, the algorithm returns “quasi-canard”, otherwise the algorithm does step six.
6. The algorithm checks whether the final point in the time series is 0.0 consumers and 3.0 resources (where the axial solution exists). If so, the algorithm returns “axial”, otherwise the algorithm returns “nothing”.

Note, the sensitivity of this algorithm to find quasi-canards can be changed by varying the top of the box in step three (changing the percentage of where the Resource isocline intersects with the Consumer axis).

**Figure S.I.4.**
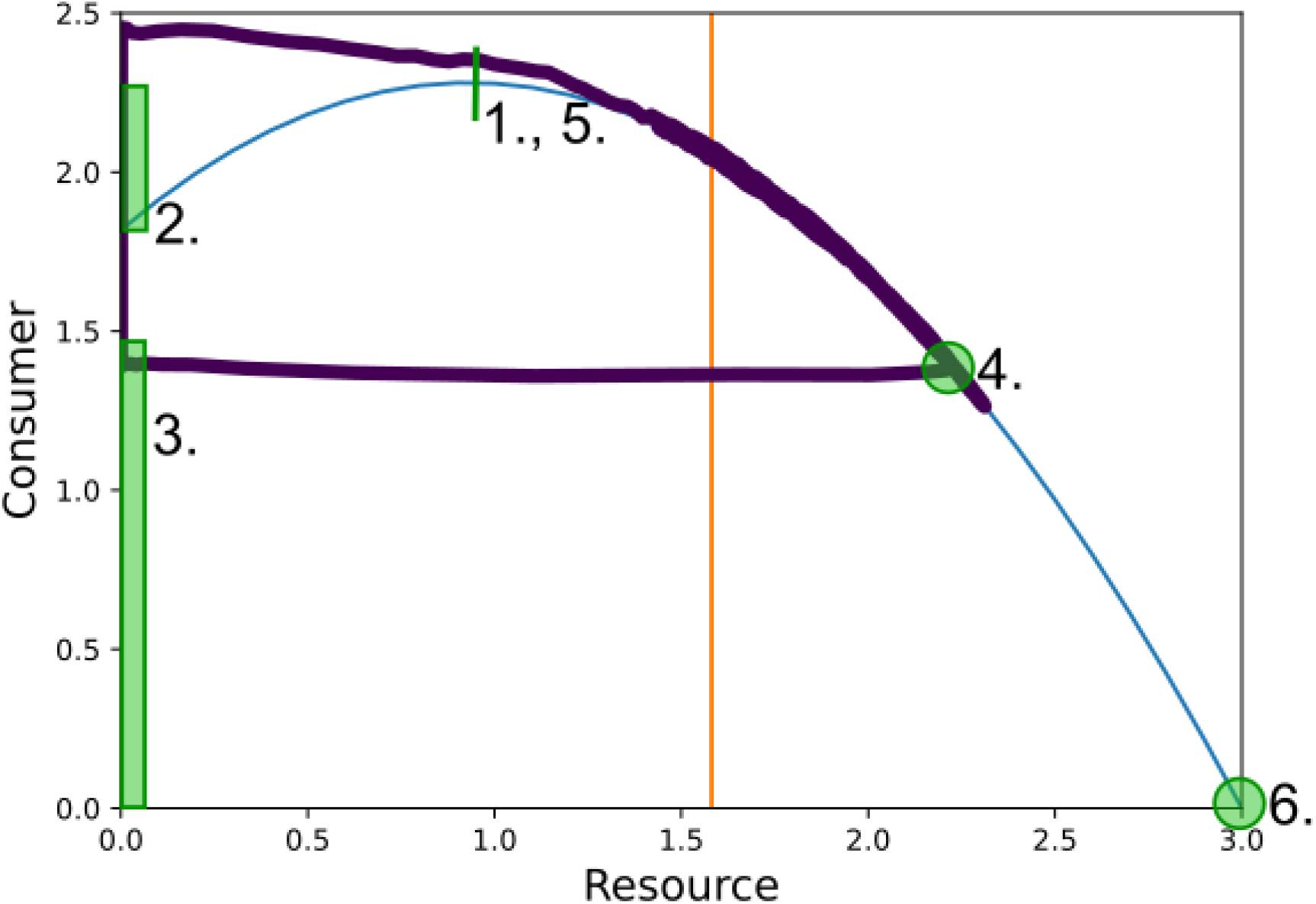
C-R phase plot with a quasi-canard, the resource isocline and the consumer isocline in purple, blue, and orange respectively. The six steps of the algorithm outlined above are depicted with green lines, boxes, and circles.

### Biologically Plausible Parameters

We used Yodzis & Innes’ [23] biologically plausible parameterization of the C-R model to test whether our sudden population disappearance results are general to other parameter sets.

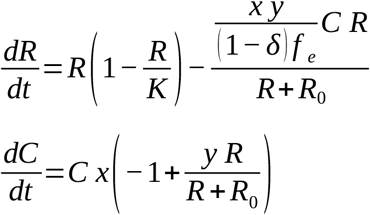

where 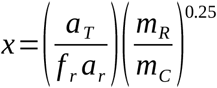

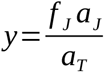

Similar to Yodzis & Innes [23], we expressed the resource body mass in terms of the mass of an equivalent endotherm operating at its physiological limit. We set our consumer as an herbivorous endotherm.

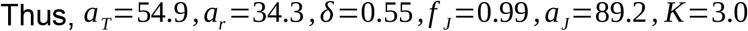

To slow the consumer relative to the resource, we multiplied the resource/consumer body mass ratio by *ε* (*ε* B = *ε m*_*ER*_/*m*_*C*_). To maintain the same biomass loss from the resource (i.e. when using the original B = *m*_*ER*_/*m*_*C*_), we set

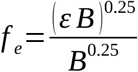

To ensure the full model was feasible (between the feasibility and Hopf boundaries in Yodzis & Innes [23]), we set B = 10^−6^ and we restricted *R*_*0*_ to [0.7,1.82].

All other methods are the same as in the main Methods section.

**Figure S.I.5.**
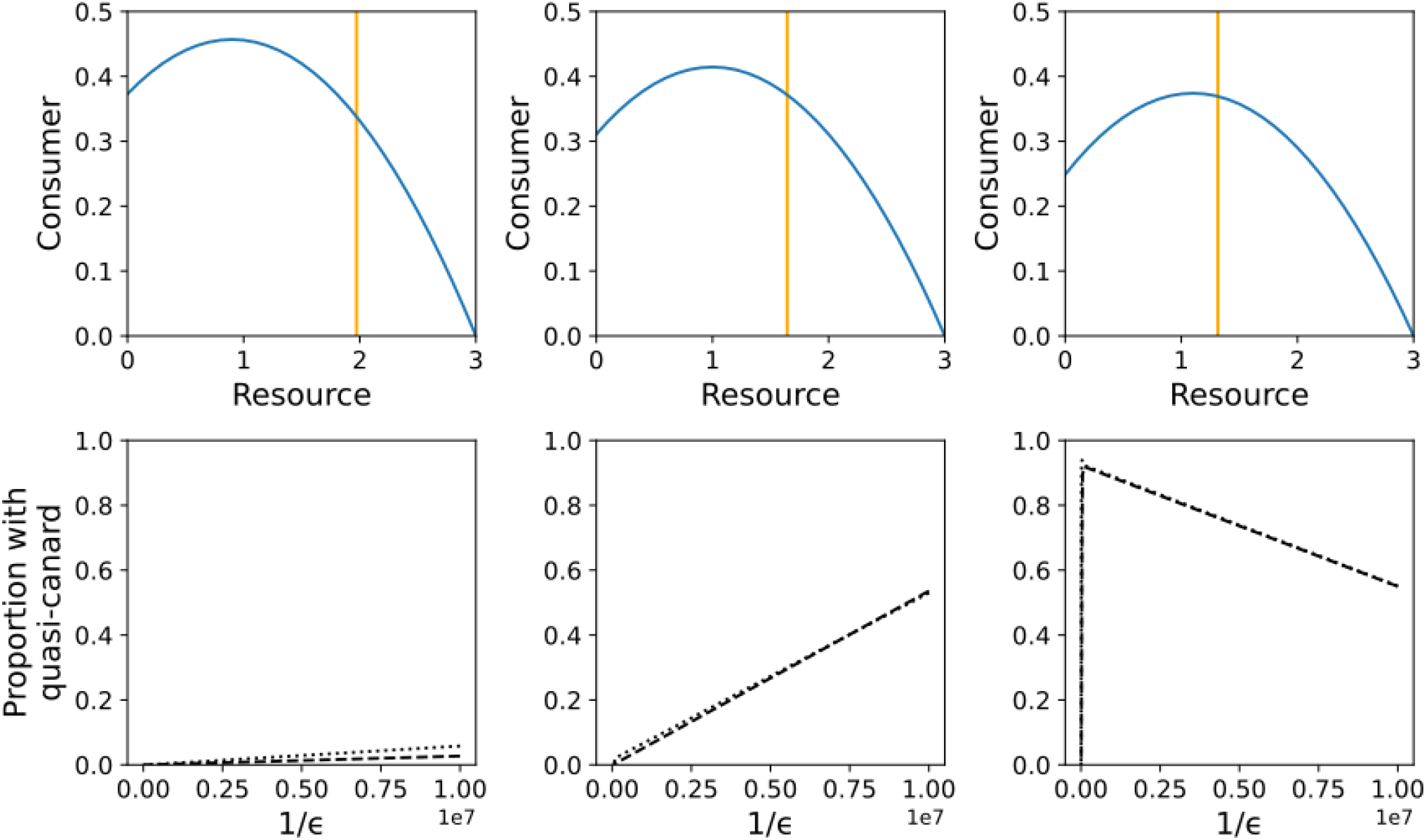
In the Yodzis & Innes (1992) model, quasi-canards can be found for a variety of efficiency values within certain boundaries of ε parameter space. a), b), c), d) resource and consumer isoclines with *R*_*0*_ values of 1.2, 1.0, 0.8 respectively. e), f), g), h) proportion of 1000 simulations per value of 1/*ε* that exhibited quasi-canards with constant *R*_*0*_ values of 1.2, 1.0, 0.8 respectively. Bold and thin dashed lines correspond to 6,000 and 24,000 time units respectively.

### Quasi-potentials

The quasi-potentials depicted below were created using the Rosenzweig-MacArthur consumer -resource model with the same parameter values as the model in the main article. We used the QPot package (version 1.2) in R to calculate the quasi-potentials (Moore *et al*. 2016). We maintained the overall intensity of noise but had different relative noise intensities between the resource and the consumer (specifically 1:4, see Moore *et al*. (2016) for how to specify different relative noise intensities).

**Figure S.I.6.**
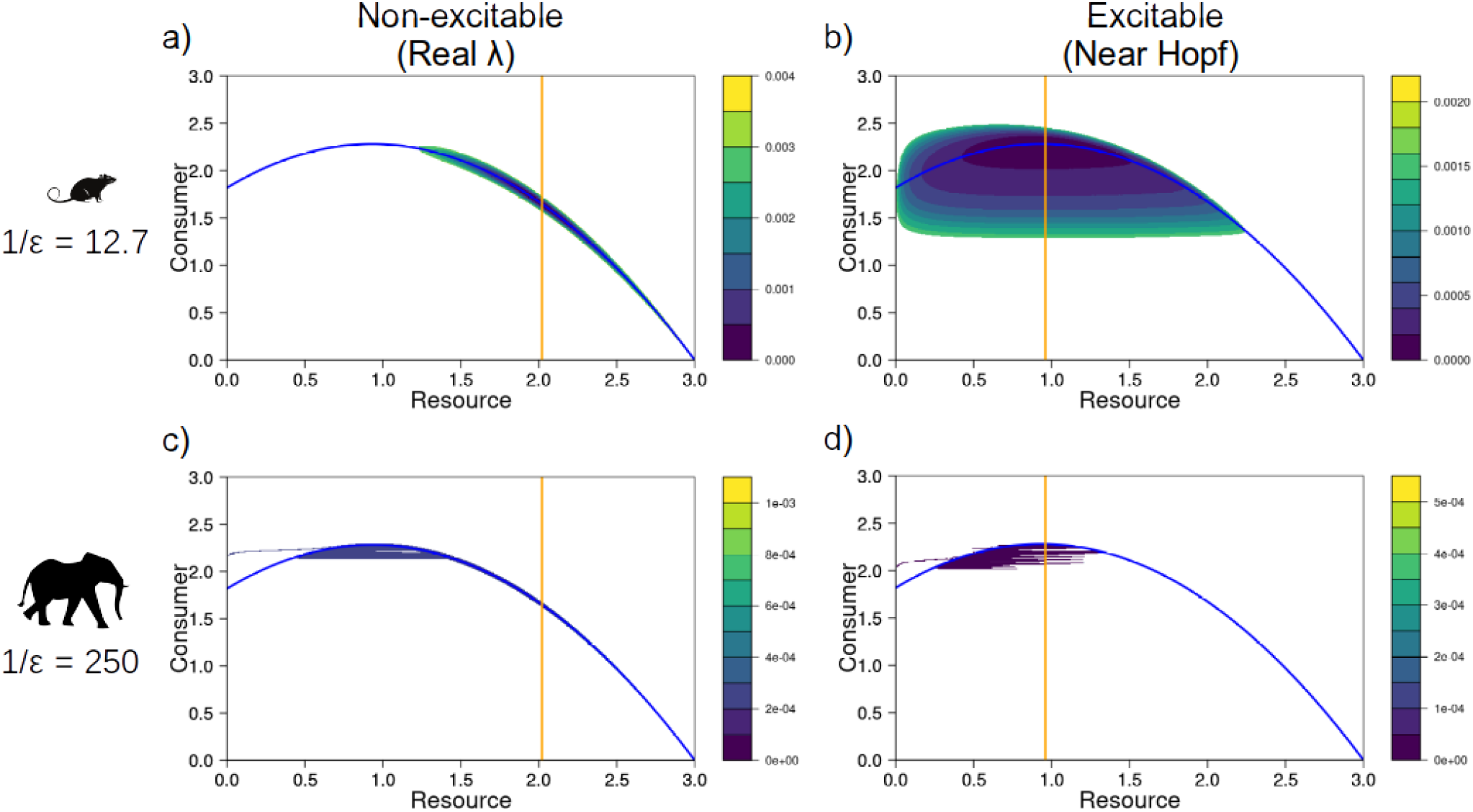
Quasi-potentials for the C-R models. The top row (a) & b)) had a 1/ε value of 12.7 and the bottom row (c) & d)) had a 1/ε value of 250. Each column of plots had efficiency values of 0.5 and 0.7 respectively. Resource and consumer isoclines are the blue and orange lines respectively.

Christopher Moore, Christopher Stieha, Ben Nolting, Maria Cameron and Karen Abbott (2016). QPot: Quasi-Potential Analysis for Stochastic Differential Equations. R package version 1.2. https://github.com/bmarkslash7/QPot.

## Notes

https://doi.org/10.5281/zenodo.5796524

### Competing Interest Statement

The authors have declared no competing interest.

### Summary of Updates

We have improved the focus of our paper and what we mean by stability. The focus of our manuscript is how consumer life history interacts with stochasticity and the deterministic skeleton (either non-excitable or excitable) to affect the stability of the consumer-resource interaction. We define stability as persistence (likelihood of not going locally extinct) and use the common theoretical and empirical metric CV (SD/mean) to define stability clearly. To help the reader with the different concepts in this manuscript, we have carefully edited our Box of Key Terms and Definitions. In our methods, we also unpack what energy flux is (a common definition of interaction strength) and how energy flux influences the consumer-resource interaction deterministically (the determinisitic skeleton sensu Higgins et al. 1997). Finally, our new Figure 1 is a clear schematic of our overall approach. Figure 1 depicts how epsilon reflects the empirical slow-fast relationship and our experiment of stochasiticity and different deterministic skeletons impacting stability (coefficient of variation) as epsilon is varied.

